# Lowest expressing microRNAs capture indispensable information: identifying cancer types

**DOI:** 10.1101/082081

**Authors:** Roni Rasnic, Nathan Linial, Michal Linial

## Abstract

The primary function of microRNAs (miRNAs) is to maintain cell homeostasis. In cancerous tissues miRNAs’ expression undergo drastic alterations. In this study, we used miRNA expression profiles from The Cancer Genome Atlas (TCGA) of 24 cancer types and 3 healthy tissues, collected from >8500 samples. We seek to classify the cancer’s origin and tissue identification using the expression from 1046 reported miRNAs. Despite an apparent uniform appearance of miRNAs among cancerous samples, we recover indispensable information from lowly expressed miRNAs regarding the cancer/tissue types. Multiclass support vector machine classification yields an average recall of 58% in identifying the correct tissue and tumor types. Data discretization has led to substantial improvement reaching an average recall of 91% (95% median). We propose a straightforward protocol as a crucial step in classifying tumors of unknown primary origin. Our counter-intuitive conclusion is that in almost all cancer types, highly expressing miRNAs mask the significant signal that lower expressed miRNAs provide.

## INTRODUCTION

Mature microRNAs (miRNAs) are short, non-coding RNA molecules. They lead to post transcriptional repression by reducing the stability of mRNAs and attenuating the translation machinery (1). In humans, there are about 1900 miRNA genes that yield ~2600 mature miRNAs. The expression levels of miRNAs in healthy tissues span 5 to 6 orders of magnitude (2).

In all multicellular organisms including humans, miRNAs were implicated in embryogenesis, tissue identity and development. As the gatekeepers of cell homeostasis (3), miRNAs respond to alteration in cell regulation that occurs during viral infection, inflammation, and numerous pathologies. In mammals, cell transformation and carcinogenesis are accompanied by drastic alterations in miRNA expression profiles (4,5). Those miRNAs that are associated with carcinogenesis and metastasis are called oncomiRs (6). They were implicated in targeting components of the cell cycle, DNA repair, oncogenes or tumor suppressor genes (7,8). Most oncomiRs are expressed in multiple cancer tissues. However, some are specific to only certain cancer tissues. For example, human miR-21 over-expression is associated with almost all cancers while miR-15 and miR-16 expressions are mostly associated with B-cell neoplasm. Other miRNAs (miR-143 and miR-145) directly regulate other oncomiRs and thus are candidates for anti-tumor therapy (9). Therefore, altered miRNA expression in cancerous versus healthy tissues is suggested as invaluable biomarkers, for cancer diagnosis and prognosis and as a lead for novel therapeutic approach (10). Despite advances in understanding cell deregulation by miRNAs in cancers for most miRNAs, a relation between expression level, diagnosis and prognosis for specific cancer types cannot be drawn (see discussion in (11,12)).

From a clinical perspective, profiling miRNAs is important for: (i) Selecting optimal treatment; (ii) Monitoring the disease’s progression; (iii) Identifying the primary origin of a metastatic cancer (12). For example, miR-10b level was shown to be a prognosis indicator for chemotherapeutic resistance in colorectal cancer (13) while miR-30c inhibits tumor chemotherapy resistance in breast cancer (14). The monitoring of minute miRNAs levels in patient’s body fluids allows a follow up for treatment and a direct assessment for disease’s progression (5,15).

Many of the samples collected from cancer patients are metastatic. Evidently, it is more challenging to treat metastatic than primary cancers (see reviews (16,17)). Importantly, the origin of up to 12-15% of all newly diagnosed cancers is unknown or uncertain (18). Due to the difficulty of identifying the origin of metastatic tissues, cancers of unknown primary origin (coined CUP) are associated with poor prognosis and inconclusive protocol for disease management (19). Consequently, there is an active line of research seeking ways to identify the type and origin of tissues from metastatic biopsies through their molecular signatures (for example (12,20,21)). Success in classifying lung squamous cell carcinomas from lung adenocarcinoma (22) was further expanded to four main subtypes of lung cancer using miRNA profiles (23). In another study, a classifier based on 47 miRNAs successfully separated 10 different tissues from ~100 samples from primary and metastatic samples (24). Expanding the task to 42 cancer types resulted in an extended set of 68 miRNAs (25) that together are used as indicators for tissue of origin in numerous biopsies from cancer patients (26). Although most measurements of miRNAs originate from cancer biopsies (27), miRNAs circulating in body fluids has become attractive candidates for early diagnosis (28,29).

The goal of our study is to use the miRNA expression profile data from The Cancer Genome Atlas (TCGA) toward the task of classifying different cancerous tissues and their disease/healthy states. TCGA provides rich molecular data from cancer patients on an unprecedented scale (30), with samples of >25 cancer types. We focused on miRNA profiles that report on the normalized expression level of each of the 1046 analyzed miRNAs. Specifically, we ask: (i) Which miRNAs best capture the information needed to distinguish the various cancerous types and origin? (ii) Which tissues are prone to false classifications? (iii) Can we find a biological interpretation of the set of informative miRNAs?

## MATERIALS AND METHODS

### Data collection

MicroRNAs (miRNAs) profiles were extracted from The Cancer Genome Atlas (TCGA, collected between October 2013 and January 2014) (31–34). We defined a class by coupling the originating tissue with the label of the sample’s state (i.e., cancerous or healthy). We limited our analysis to classes with over 50 samples. In total we collected 27 classes. Three classes of healthy tissues are abbreviated as Kidney, Thyroid and Breast. Additionally, 24 cancerous classes and their acronyms are listed: adrenocortical carcinoma (ACC), bladder urothelial carcinoma (BLCA), brain lower grade glioma (LGG), breast invasive carcinoma (BRCA), cervical squamous cell carcinoma and endocervical adenocarcinoma (CESC), colon adenocarcinoma (COAD), esophageal carcinoma (ESCA), head and neck squamous cell carcinoma (HNSC), kidney chromophobe (KICH), kidney renal clear cell carcinoma (KIRC), kidney renal papillary cell carcinoma (KIRP), liver hepatocellular carcinoma (LIHC), lung adenocarcinoma (LUAD), lung squamous cell carcinoma (LUSC), pancreatic adenocarcinoma (PAAD), pheochromocytoma and paraganglioma (PCPG), prostate adenocarcinoma (PRAD), rectum adenocarcinoma (READ), sarcoma (SARC), skin cutaneous melanoma (SKCM), stomach adenocarcinoma (STAD), thyroid carcinoma (THCA), uterine carcinosarcoma (UCS) and uterine corpus endometrial carcinoma (UCEC).

The 27 classes include 8522 distinct samples. Each sample is associated with the expression of 1046 miRNAs. We assessed the similarity among patient samples by calculating the cosine of the angle between the vectors of miRNA expression. These cosines were used as a proxy to the initial separation ability and similarity among classes. The lists of all samples and miRNAs’ TCGA identifiers are provided in supplementary data Table S1.

### Data transformation

We apply a rough discretization to the data, according to miRNA expression level, measured in reads per million (RPM). We start with a dichotomy - one category for expression levels strictly between 0 and 30 RPM, and a second category for everything else. A somewhat more refined classification is a trichotomy, that partitions the data to non-expressed, lowly expressed (expression levels from 0 to 30 RPM) and highly expressed (>30 RPM). On average, each sample has 306 lowly expressed miRNAs (s.d. of 43.91), and 163 highly expressed miRNAs (s.d. of 18.58).

### Machine learning model

Human TCGA samples were classified to the 27 classes (see above) by training a multiclass support vector machine (SVM) classifier (35). We used a Python implementation based on the scikit-learn package (36). The training sets for the raw data, the dichotomy and the trichotomy were limited to 40% of the samples of each class. The sensitivity of the SVM classifier to the size of the training set was tested. The tests were repeated with training set comprised of 20% to 80% of each class of samples, with increments of 10%. Our results for both data transformations were consistent and stable for training sets of at least 40% of the samples and somewhat less stable for training sets below the 40% threshold. Both dichotomy and trichotomy schemes were run 1000-10,000 times each. Scikit-learn cross-validation mechanism was used in order to prevent over-fitting.

Prior to data analysis we randomly partition all data into 40% that are unseen and 60% that are seen. We create 5 training sets each of which is comprised of a random subset of two-thirds of seen part. For each set, two classification matrices were created, one based on the 30 RPM threshold, and the other based on a 60 RPM threshold, for separating the lowly-expressed from the highly-expressed. We then run our machine-learning procedure on each of these 5 sets and test them on the unseen part of the data. Our final classification is chosen by carrying a plurality vote among these 10 classification results.

### Performance evaluation and error analysis

We assessed our models according to their average recall rate for each of the classes. Recall is defined as [true positive] / [true positive + false negative]. The average, median and standard deviation (s.d.) were measured for over 300 independent runs of the SVM protocol on raw or discretized data.

We carried out two types of error analysis: At the sample level and at the class level. First, we analyzed the errors made per each of the samples over distinct runs (Sample-based Error). A second error analysis concerns a class misclassifications and the identity of the faulty assignment (Class-based Error). The plurality vote was implemented in our final classification in order to overcome Sample-based Errors. We have merged the classes READ and COAD in order to reduce the extent of Class-based Errors.

### Informative miRNA extraction

Several methodologies were applied to identify and select a minimal set of miRNAs that are most informative towards a successful classification task. Our best results were obtained using the above described trichotomy. Specifically, for each miRNA we create an 81-dimensional vector, with 3 coordinates for each of the 27 classes (24 cancer types and 3 healthy tissues). These three coordinates are defined as follows: If there are N samples in this class for the miRNA at hand, of which N_0 are not expressed, N_1 are between 0 and 30 RPM and N_2 are highly expressed, then we set the first coordinate at N_0/N, the second to N_1/N and the third to N_2/N. We then calculate the standard deviation (s.d.) of each of the expression levels among the 27 classes separately, and calculate the sum of the s.d. Formally:

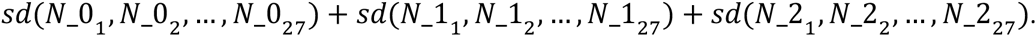

We expect miRNAs of higher summed s.d. values to be more informative, since the s.d. captures the miRNA’s variability among different classes. We chose not to assign distinct weights to different classes despite their differing samples’ size (ranging from 57 for Uterine Carcinosarcoma to 1066 for Breast Invasive Carcinoma).

## RESULTS

### Many cancerous tissues have similar miRNA profiles

The main goal of our study is to classify cancer classes using the information encoded by miRNA profiles from patients. For most cancer types, a small number of highly expressed miRNAs undergo a drastic change in expression level along the progression of the disease. Therefore, it is a commonly held view that uniquely expressed miRNAs might be used for diagnosis, prognosis and possibly also for disease treatment. Here we revisit this view and test the miRNA heterogeneity and expression consistency in an unbiased way. We analyze many different cancer types and tissues, and thousands of patient samples.

We wanted to get a sense of the similarity between the vectors of miRNA reads among samples from each given tissue. We found high correlations among the miRNA expression vectors when both samples are healthy or when both are diseased (within the same tissue). The correlation is much lower for healthy vs. cancerous one (data not shown). Figure 1 shows an analysis for all pairs of patients with lung adenocarcinoma samples (521 samples, abbreviated LUAD) and matched healthy samples (~50 samples). The similarity between each pair is indicated (cosine values, see Materials and Methods). We show that healthy and cancerous tissues exhibit different patterns. Importantly, healthy samples are similar to each another as are the diseased samples.

**Figure 1.**
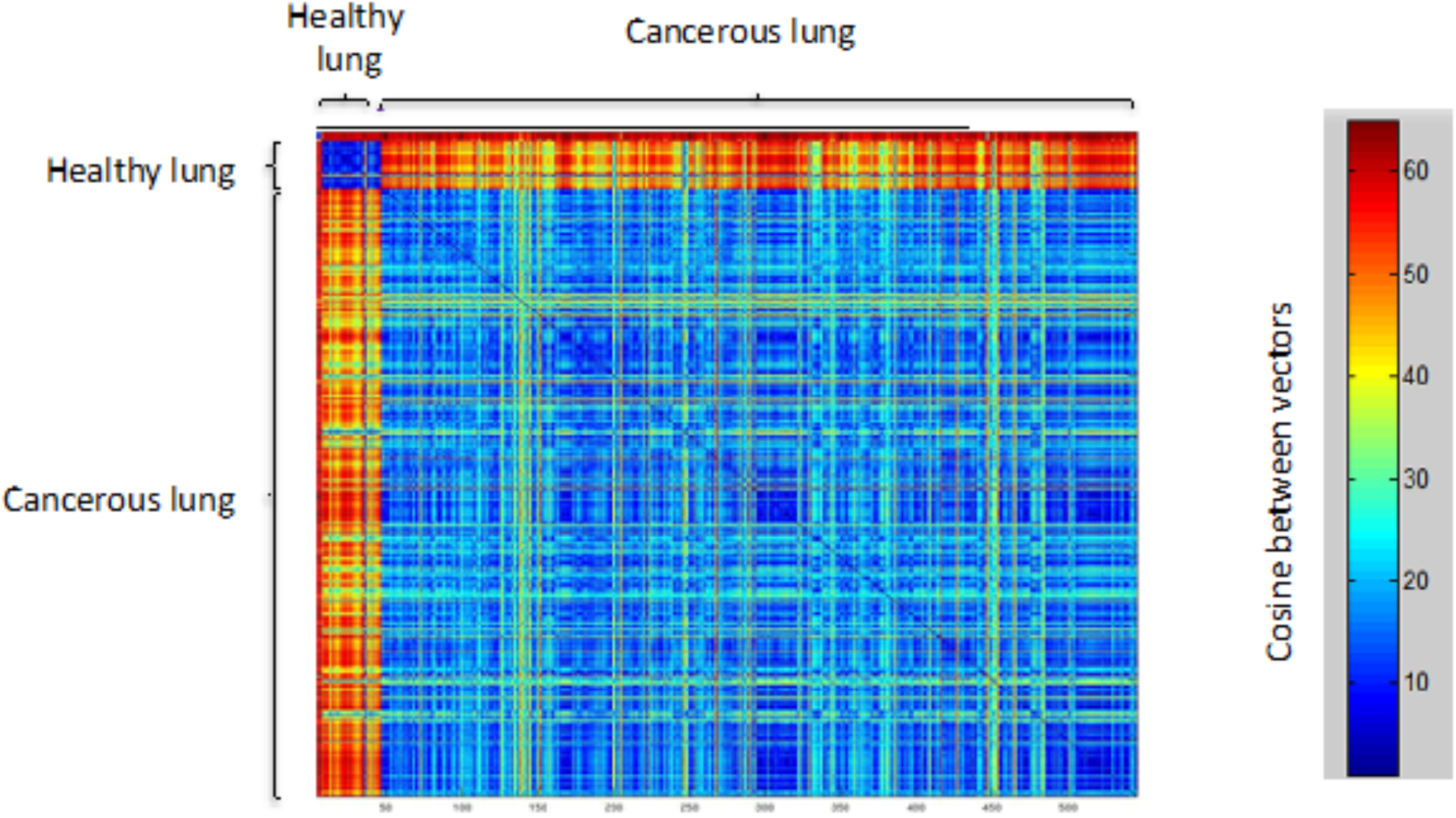
Distance matrix of the relations between lung adenocarcinoma patients’ samples and healthy lung samples. There are 50 healthy samples and 521 diseased samples. The distance is the cosine between each pair of samples. The larger the angle, the larger is the distance between the paired miRNA expression vectors. Blue to red indicating the range from similarity to maximal distance.

We tested whether this property (Figure 1) extends to all other cancerous and healthy tissues. We therefore restricted our analysis to tissues for which samples from both healthy and cancerous patients are available. As Figure 2A shows, each healthy tissue is well characterized by its miRNAs’ profile. For example, breast tissues in different patients have very similar miRNA profiles, and differ significantly from liver profiles. Even the two types of lung tissues are visibly distinguishable from each other. However, the distance matrix for the diseased samples (5 classes, ~2700 patients, Figure 2B) is relatively uniform. While all the diseased tissues are rather similar using our similarity matrix, a somewhat higher similarity is visible between the two lung diseases, the adenocarcinoma (LUAD) and lung squamous cell carcinoma (LUSC). We concluded that the trend that we observed in Figure 1 applies to other tissues. Namely, the distance among all pairs of diseased tissues is lower than that among the corresponding healthy tissues.

**Figure 2.**
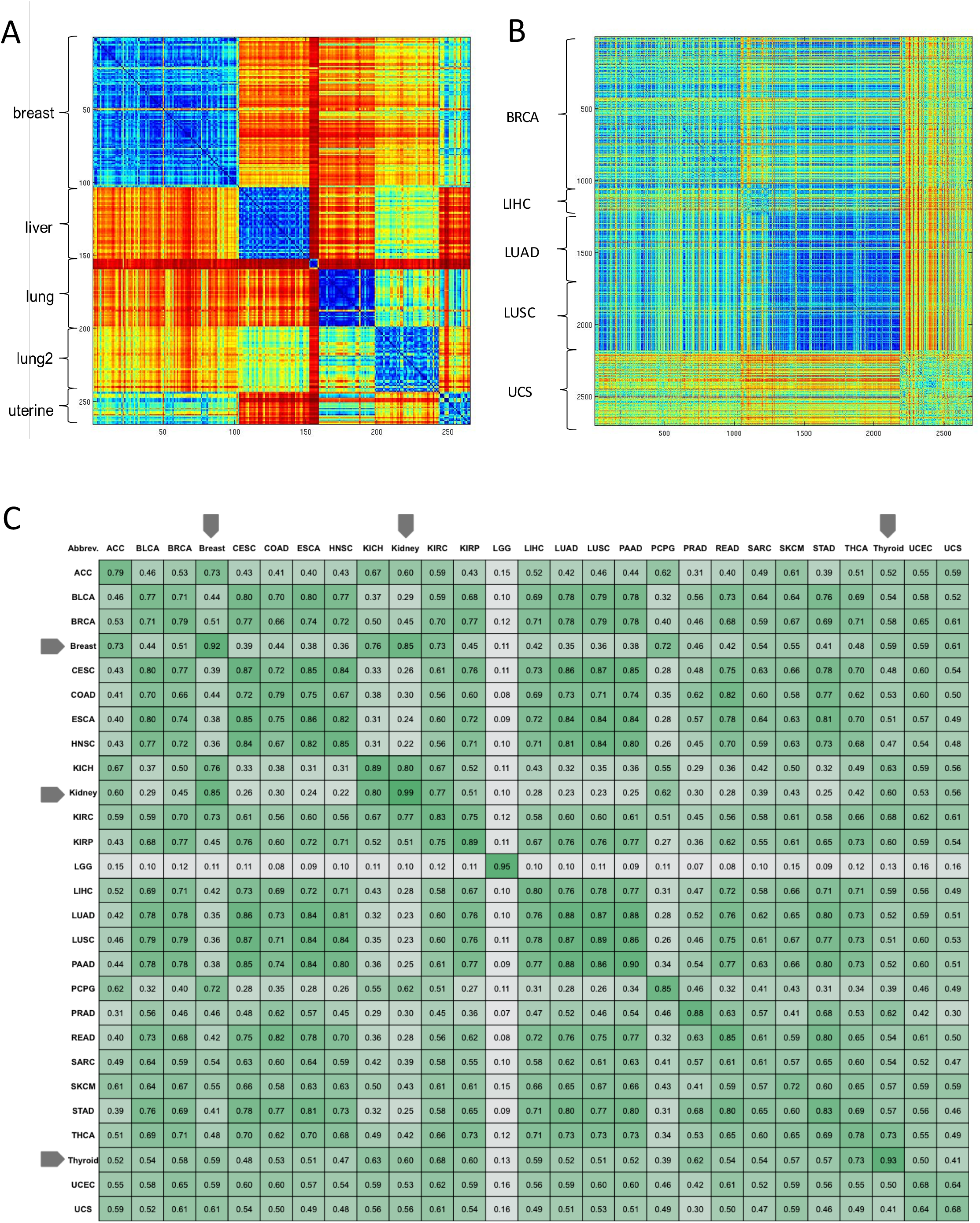
Analysis of the relations between samples from different tissue origin. **(A)** Distance matrix of healthy samples from 5 sources. The healthy samples are far more similar within the tissue than among tissues, substantial similarity is observed between healthy samples from lung adenocarcinoma (LUAD) and lung squamous cell carcinoma (LUSC) that are marked as lung and lung2, respectively. **(B)** Matched diseased samples from the same tissues reported in (A). The number of patients from each class is indicated (axis Y). Recall that for the machine learning approach we excluded classes that are supported by <50 samples (see Materials and Methods). As oppose to the healthy samples, the distances between the patient samples are rather uniform with no clear partition between diseased classes. Blue to red indicating the range from similarity to maximal distance. **(C)** A ‘bird view’ of the miRNA-based information for >8500 TCGA samples. All 27 classes are listed in an alphabetical order. The classes that are associated with healthy samples (Breast, Kidney and Thyroid) are marked by arrows. Note that the correlation within the same class (the diagonal) is not necessarily maximal (e.g., 0.61 for Sarcoma, SARC). The brain lower grade glioma (LGG) is an outlier with maximal distance to any other class.

Figure 2C offers a bird's eye view of miRNA-based information for all 8522 samples that were included in our analysis, according to 27 classes discussed in this study. The entries show the average of the correlations over all samples in each pair of classes. Note that the values on the diagonal are only slightly higher than the rest of the values in this symmetric matrix. It is noteworthy that that brain lower grade glioma (LGG) markedly differs from other cancerous tissues, yet exhibiting a significant self-similarity. Additionally, sarcoma samples (SARC, diagonal, Figure 2C) exhibit rather low (0.61) average correlation, suggesting high variability of the miRNA expression in this cancer type.

### Lowest expressed miRNAs provide differentiating information

In view of the relatively homogenous pattern of miRNA profiles among diseased tissues (Figure 1), it is clear that the identification of cancer classes cannot be trivially based on a predefined simple set of miRNAs. Thus, we sought a classifier that can correctly label a tissue of origin based on information that is globally captured by the miRNAs’ profiles. To this end, we applied a multiclass support vector machine (SVM) classifier with a randomly chosen training sets (Figure 3A). The flow of the methodologies used in this study is shown. Our classifier employs two variations of data transformation, following by a step that benefits from studying the typical errors in our classification task (Figure 3A).

**Figure 3.**
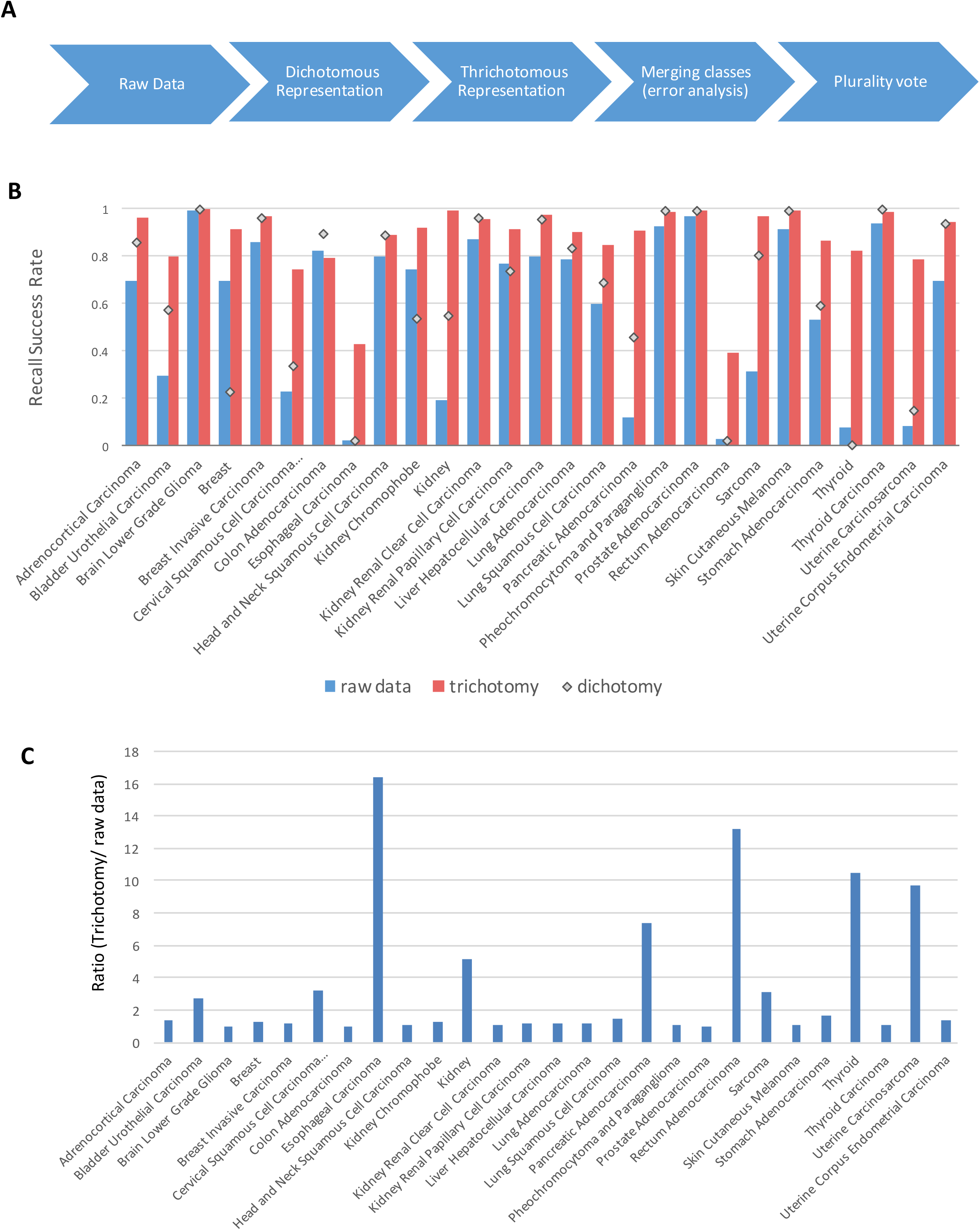
Classification results following data transformation. **(A)** A scheme of the workflow. Each knob described either a data transformation or a refinement of the used method. **(B)** Average success rates for each class while using raw data (blue), data in a dichotomous representation (empty diamonds) and data following a trichotomy representation (red). **(C)** Fold improvement achieved by applying trichotomy transformation with respect to the results from raw data. The classes are sorted alphabetically.

SVM applied to the raw data to identify the tissue of origin achieves an average success rate of 58%, and a median of 69% (27 classes) (Figure 3B, blue). Note that naïve guessingwould yield ~4% success rate for classifying a sample to any of the 27 classes. Interestingly, for 10 out of the 27 classes attains recall rate higher than 80%, and for 5 classes the success rate is even above 90%. However, the prediction for some classes is very poor (<10%). The best performance is associated with LGG, as could be anticipated from the observation in Figure 2C.

However, for clinical purposes, a success rate of 58% is unsatisfactory. We thus focused on searching a subset of miRNAs that are most informative towards the task of classifying tissue of origin and sample status (diseases, healthy). The expression levels of miRNA exhibit a high dynamic range of 5-6 order of magnitudes and expression is expressed in RPM (reads per millions). Furthermore, it turns out that only a small number of miRNAs (20–25) account for the vast majority of the reads in the samples (96-98%, not shown). Based on these observation, we tested the potential of discretizing miRNAs’ raw values, thus allowing a representation that emphasizes the contribution of lowly expressed miRNAs. We anticipated that the long tail of miRNAs might be more informative for the classification purposes with respect to a small set of dominating miRNAs.

We applied a simple dichotomy that partitions the lowly expressed miRNA (< 30 RPM) vs. the rest of the miRNAs. Typically, ~70% of the miRNAs are expressed at the range of >0 to 30 RPM. This simple transformation (see Materials and Methods) dramatically reduces the volume of data, and concentrates on only ~ 0.02% of the original signal (i.e., the sum total of miRNAs expression values). Importantly, the application of naïve dichotomy already improves the performance of our classifier to an average success rate of 66% (median of 80%, Figure 3B).

We sought additional data processing procedures to further improve the classification performance. In dichotomy we treat equally miRNAs that are expressed at levels above the threshold (>30 RPM) and those that are not expressed at all in a specific class. We next separate these two sets, and replace the data transformation to a trichotomy (Figure 3A, see Materials and Methods). This transformation yields a substantial improvement. The classifier thus reaches an average success rate of 88% (median 91%, Figure 3B and Table1). Figure 3C shows the improvement in the classification success for the trichotomy mode with respect to the original raw data. It is clear that some classes benefit more from this transformation than others. For 8 out of the27 classes the observed improvement was >3 folds (Figure 3C).

**Table 1.**
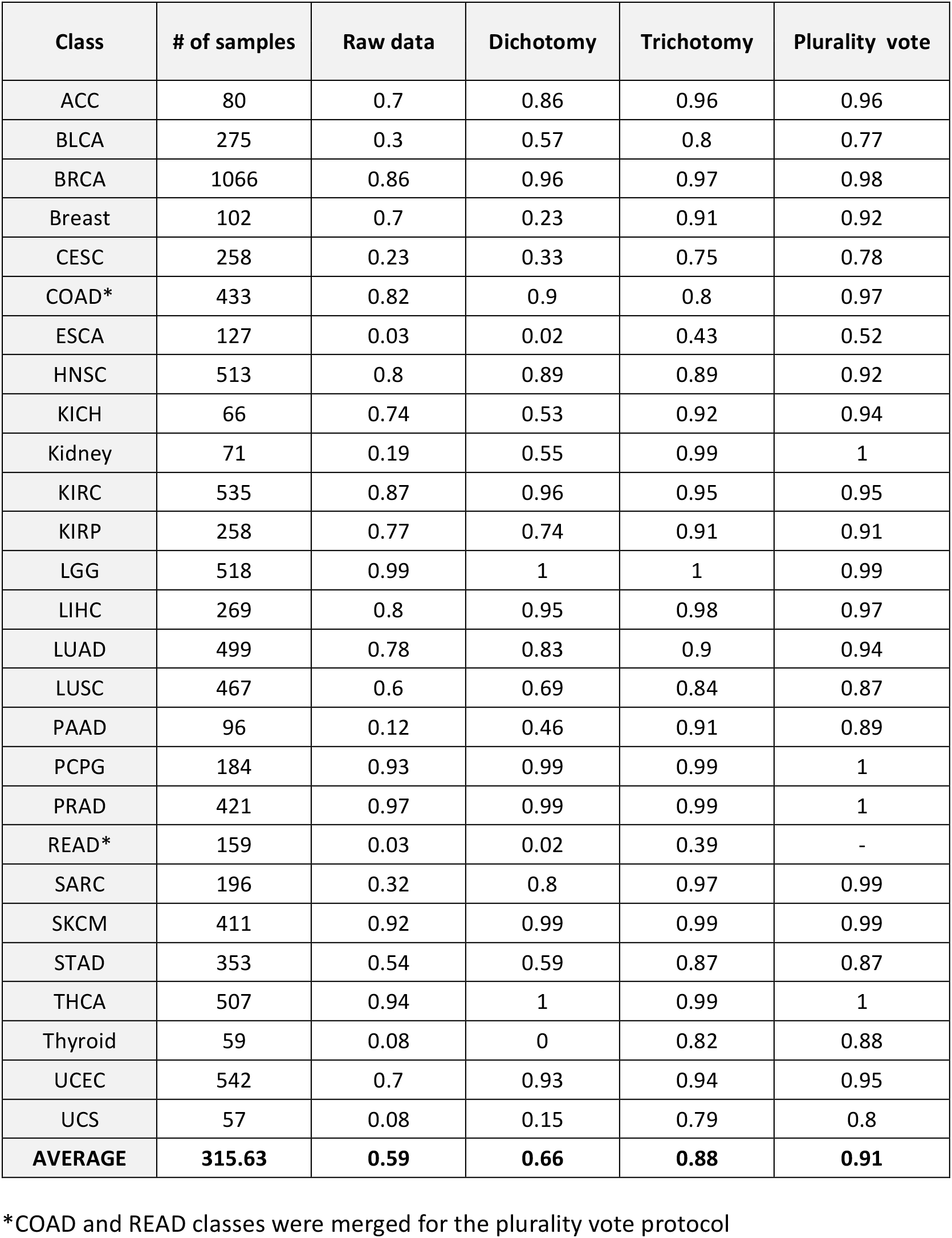
Average classification success for classification of the 27 classes by various data transformation protocols.

We analyzed the dependence of the success rate on the chosen threshold (<30 RPM) and observed that the recall rate remained virtually unchanged for thresholds ranging between 5 to 300 RPM. However, raising the threshold above 300 RPM affects the performance negatively, suggesting that at this range of parameter we are actually adding noisy information (Figure 4). It is evident that for both the dichotomy and trichotomy modes, information within the very low expression levels is critical for the classification task. The contribution of the very lowly-expressed miRNAs to the classification task was further assessed. We zoomed on miRNAs expressed at levels x, where x ranges from >0 to 30. We trained the classifier again for different values of x and artificially relabeled miRNAs that are smaller than x as non-expressed. When marking as non-expressed miRNAs with expression levels under 10 RPM, there was no significant change to the classification results, while applying x > 10 RPM led to a decline in performance. Typically, only ~8% of the miRNAs are listed in the range of 10-30 RPM, while 25% of the entire list of miRNAs are within the range of 1 to 10 RPM. We conclude that lowly expressed miRNAs that carry fundamental information for the classifier, mostly (but not entirely) reside in the 10 to 30 RPM range.

**Figure 4.**
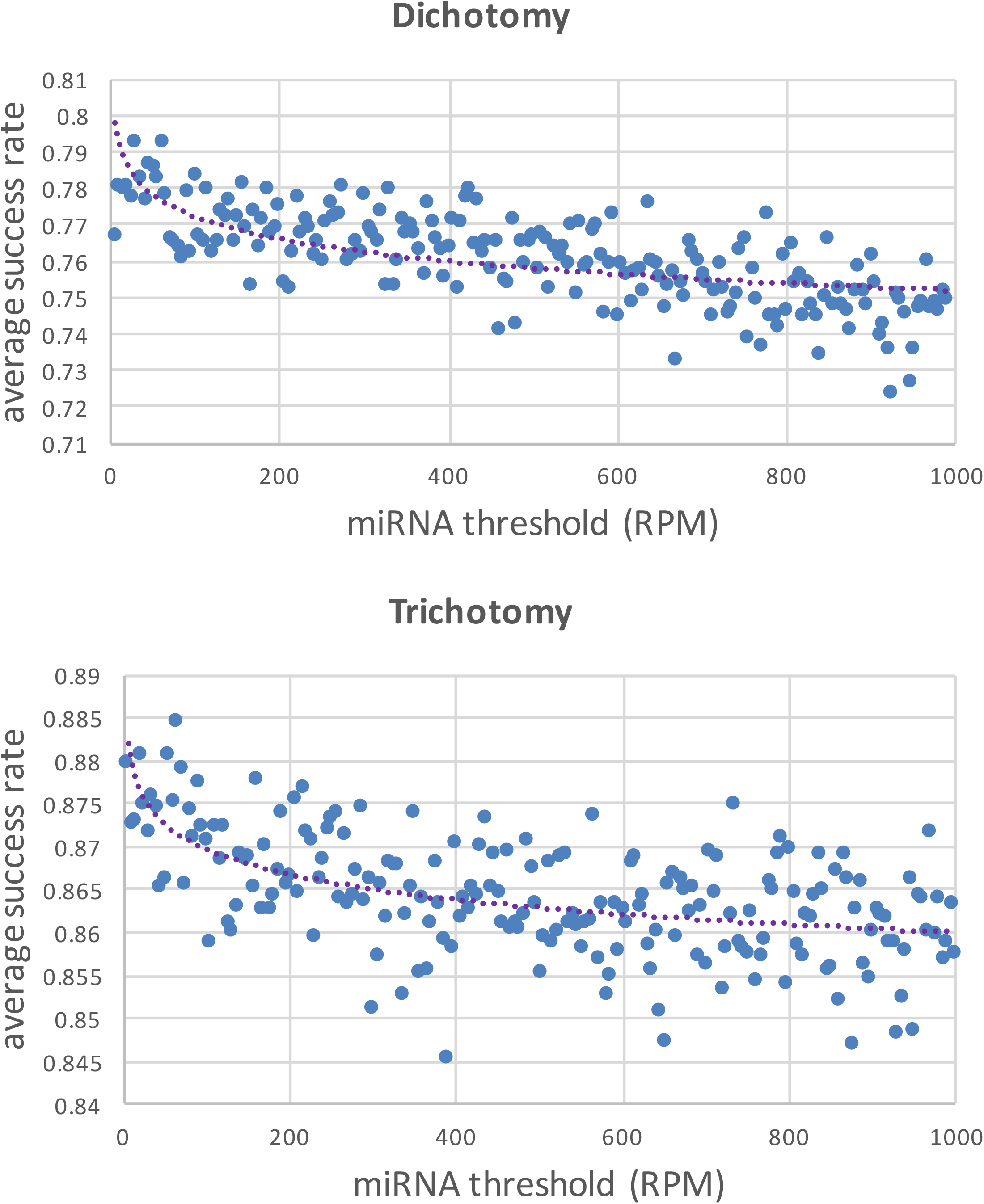
Average success rate when applying 200 separation thresholds from 5 to 1000 RPM with 5 RPM increments. The average success rate obtained for the dichotomous transformation and the trichotomous transformations are on Top and Bottom, respectively. The drop in performance supports the notion of increasing noisy data by increasing the threshold. Note a difference in the average performance for dichotomy vs. trichotomy transformations.

### Not all tissue types are equally hard to classify

As mentioned, trichotomy has resulted in a substantial improvement for many classified tissues. For 11 out of the 27 classes, we consistently reached a success rate of 95% or above (Table 1). Importantly, the classification success does not immediately reflect the sample size. For example, a 96% success for Adrenocortical Carcinoma (ACC) is based on only 80 samples while lung squamous cell carcinoma (LUSC) achieved only 84% success with a sample size which is 6-fold larger (Table 1). Specifically, we are doing poorly on classifying esophageal carcinoma (ESCA) and rectum adenocarcinoma (READ), with a success rate of 43% and 39%, respectively.

We tested the sensitivity of the classification performance with respect to alternative machine learning methods: Random Forest and SVM with polynomial and radial basis function (RBF) kernels. The results for Random Forest and SVM with polynomial kernel were somewhat worse than the used SVM (see Materials and Methods). RBF kernel gave an improved performance on the raw data, but failed to improve following data discretization and data transformation.

### A third of the classification errors are interpretable

We tried to refine our understanding on features that govern the success / failure in the classification task. To this end we inspected the results in view of the different errors that are made. Figure 5 presents a “wheel view” for all 27 classes with a visual indication on the samples’ size for each class, and the typical errors. Cross-edges in the graph and their color indicate the extent and nature of all misclassifications. The wheel indicates the true-positive and false-positive (inner arc), as well as the true-positives and false-negatives (middle arc). From the outermost arc one can estimate the sum of the two error types. The source data is available in Supplemental Table S2.

**Figure 5.**
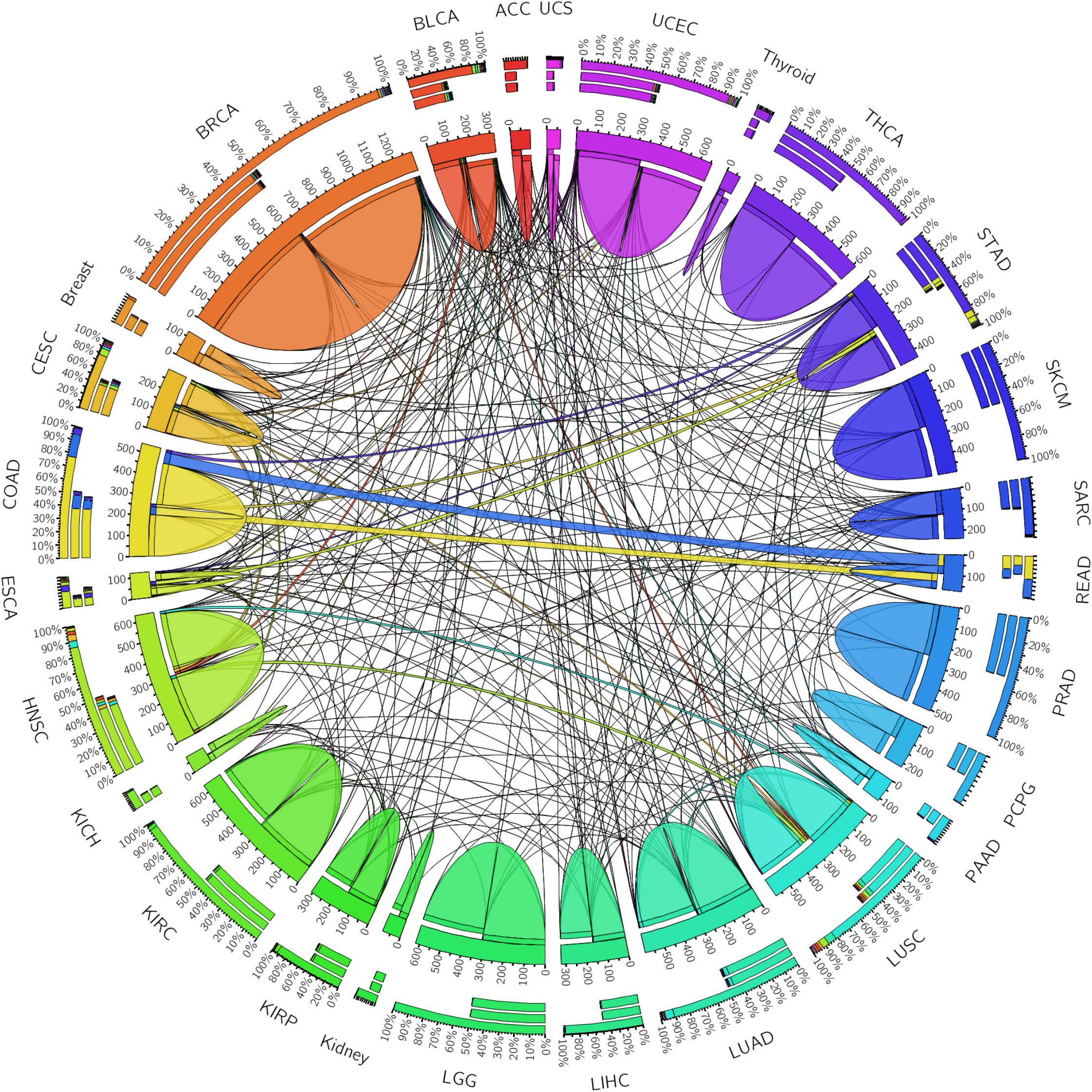
A circular view including 27 classes according to the error types and the size of each class. Each class is associated with two quantifiers and their summation. The inner arc displays the distribution of the classes classified as the subjected class (i.e., true-positive and false-positive), the middle arc displays the distribution of classification of the samples which belong to the subjected class (i.e., true-positive and false-negative). The outer arc is the summation of the inner and middle arcs. The number of edges and their color-coded display the extent and nature of all missed classifications. Note the two-sided errors for the classes of COAD and READ. Each class is color coded and the errors are colored by the origin of the misclassified classes. Supplementary data Table S2 depicts the average classification results per class for 100 SVM runs. Table 2 is a visualization of the data presents in Table S2.

We are often unable to interpret the errors in classification. We categorized the errors by biological relevance of the classification errors: (a) Cancerous state errors: Correct classified tissue but failure in determining healthy vs. diseased state. These are restricted to the 3 instances of tissues that are represented by both healthy and cancerous instances. (b) Intra-tissue errors: misclassification of one class to another that is anatomically adjacent. Such errors are limited to classes with multiple diseases of the same/ related organ. (c) Inter-class errors: classification mistakes that are not obviously interpretable. Figure 6A shows schematically the possible error types for all 27 classes. Figure 6B indicates the distribution of different error types among the relevant categories of errors. Figure 6C focuses on an example for inter-class error and the source for false positives. It shows that in the case of BLCA, the majority of the false positives (10.6% out of 12.7%) derive from misclassification from 6 classes of cancer carcinomas (additional classes, that contribute <1% of false positives each are not shown).

**Figure 6.**
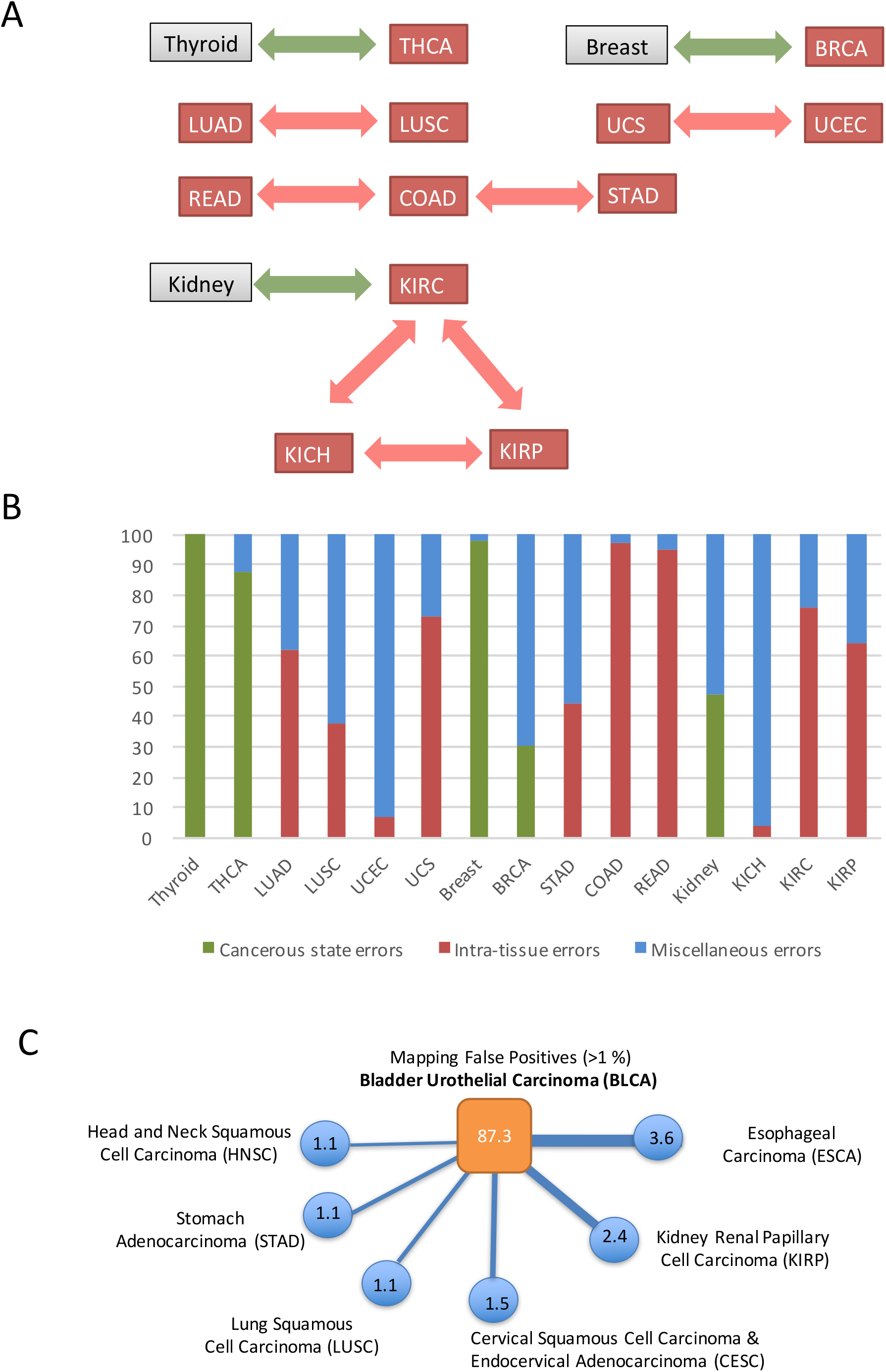
Analysis of classification errors from similar sources. **(A)** A diagram of the affiliated tissues. Healthy and cancerous tissues derived from the same tissues are connected with a green arrow. Adjacent cancerous tissues are connected with a red arrow to reflect the anatomical relatedness. **(B)** The rate of intra-tissue and cancerous state tissue out of the total errors for each of the relevant classes. (C) An example for the source of misclassification focusing on the false positives. The fraction of false positives occupies 12.7%. The main classes that contribute to the false positive are shown along their contribution. The indicated 6 classes shown account for 85% of all false positives for this class (BLCA).

### Improving classification success from errors’ consistency

As Figures 5-6 demonstrate, many errors are recurrent among specific classes. We tested the effect of merging classes that are characterized by abundant, bi-directional false assignments by the classifier. The most prominent case involves Adenocarcinoma of the Rectum and Colon (READ and COAD, respectively). Merging these two classes has improved the average recall rate, while merging other pairs of classes has worsened the overall classification success (not shown).

When we examined the errors per sample it transpired that for certain samples, classification varied with the (random) choice of training sets. A plurality vote protocol was applied in order to reduce this dependency (see Materials and Methods). The plurality vote led to an additional improvement in the average classification success. For most classes the improvement is rather modest. However, merging the Rectum (READ) and Colon Adenocarcinoma (COAD) classes that were reported with 39% and 80% success, respectively was highly beneficial. Actually the merged collection of samples reached 97% recall. The average recall rate for the valid 26 classes has reached a further improvement to 91%, with a median of 95% (Table 1).

Additional tests in which the classifier was trained to identify the clinical stages of the disease (i.e., cancer stage 1 versus stage 3) and other global features, like age and gender, resulted in poor performance and will not be further discussed.

### Lowest expressing miRNA unveil informative miRNAs

Notwithstanding the impressive success rate of the classifier (91%, with 95% median), we are still unable to quantify the contribution of any specific miRNA to this success. To this end, we designed a ranking score for miRNAs according to their potential in contributing to the classifier success (see Materials and Methods). Empirically, we compared the results achieved when classifying with a reduced vector of miRNA expression. We randomly drew various groups of 50 miRNAs out of the 1046 miRNA discussed in this study (Table S1) and tested the classification results when using the data as is (raw data), as well as following the trichotomy transformation (Figure 7A). We also tested the results when selecting the 50 most informative miRNA (according to rank by sum of the statistical variation, see Materials and Methods). We repeated this randomization test for a growing set of miRNAs. Note that as the set size grows, a random set tends to include more informative miRNAs (e.g., for 200 miRNAs). Consequently, the differences between the performance of the randomized set and the most informative set is reduced. Already the 50 most informative miRNAs attain an impressive 75% success rate, suggesting that much of the classification signal is already captured by the set that is characterized by maximal variance (Figure 7A). With 300 of the most informative miRNAs, success rate reached on average 87%, similar to what is achieved when all the miRNAs are employed along with the trichotomy transformation.

**Figure 7.**
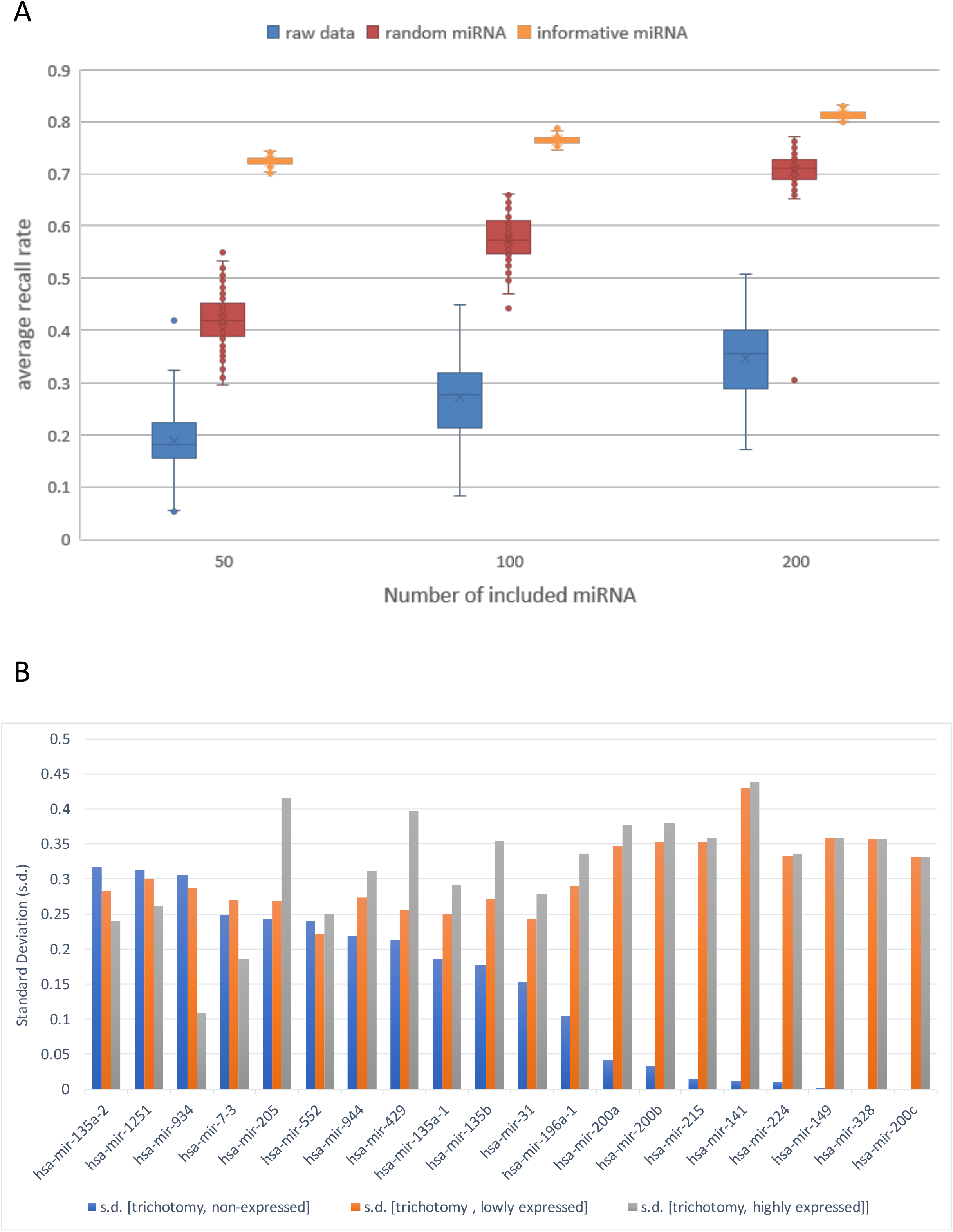
Average success rate achieved when limiting the number of miRNAs. **(A)** The performance of the classification scheme following reduction of the miRNA sample size to 50, 100 and 200 is shown. The blue markers show the results with randomly selected miRNAs in their raw form are used. The red markers were run with randomly selected miRNA in the trichotomy format. The orange markers were run with the top informative miRNA in the trichotomy form (see Materials and Methods). The box-plot presents 75% of the data. The s.d. for each column for the raw data were 6.07%, 7.59% and 7.19% (for 50,100 and 200 miRNAs accordingly). The s.d. for each column forthe random miRNA in the trichotomy representation were 5.09%, 4.15% and 4.8%. The s.d. for each of the informative trichotomy runs were 0.01%. **(B)** A list of 20 most informative miRNAs according to the s.d. of the three vectors used to define the trichotomy transformation. The miRNAs are sorted according to the s.d. for the non-expressed vector. The full list of miRNAs associated with the s.d. for each of the three coordinates and ranked by the variability sum is available in Supplemental Table S3.

We tested whether the miRNAs that contribute most to the classification task are indeed those that were implicated in cancer progression, prognosis, and diagnosis. To this end, we analyzedthe 20 most informative miRNAs by the criterion used in Figure 7A. The source of variability is shown as reflected by the s.d. associated with the 3 coordinates for each class (no expression, lowly expressed (0<x<30) and highly expressed (>30 RPM). For a complete rank of informative miRNAs see Supplemental Table S3. It is clear that some informative miRNAs rely on variability in the vector of highly expressed (Figure 7B, compare miR-205 to miR 934). For others, no variability is observed in the vector for non-expressed (Figure 7B, blue). While both hsa-mir-141 and hsa-mir-215 are among the most informative miRNAs, their source of variability is rather different. While hsa-miR-205 is characterized by a very large variation to in the highly expressed section, yet a substantial variability is recorded for this miRNA also in the other two categories (non-expressed and lowly expression).

## Discussion

### Cellular quantities of miRNAs

Our results uncover the power of the least expressed miRNAs in distinguishing a large collection of tissues and cancerous types. The common notion is that the quantity of mature miRNAs in the cell is critical for cell regulation. Specifically, excess of a specific type of miRNA potentially titrates out genuine mRNA targets (37). As a secondary effect, an overflow of specific miRNA will most likely result in its binding to non-genuine targets, the so-called called “off target” effect (38). The overall quantitative effect of miRNA levels is discussed in terms of competition (39,40) and cooperativity (41). Most studies that consider miRNAs in living cells tend to be biased toward the analysis of highly expressed miRNAs. Within such framework, lowly expressed miRNAs are most likely discarded with the premise that they have no impact of the mass effect.

In this study, we show for the first time that for the task of cancer classification, miRNA-based information is encoded by the minute expression of miRNAs that together account for only 0.02% of the total reads. Moreover, it is apparently the long distributional tail of miRNA expression that carries the signal for correctly identifying multiple cancer / tissue types. We actually show that including the entire profile of miRNAs leads to poorer performance as the principal, informative signal for class identification is masked.

A practical consequence of our observation concerns the preferred sequencing depth needed for a routine miRNA profiling. The high coverage that is reported for all samples in TCGA, entails a high sensitivity. Specifically, about 45-50% of the miRNAs, in almost all classes are expressed at a level of 0 to 1 RPM. The miRNA list with values of 1-10 RPM accounts for an additional 25% of the miRNA list. Classically, when sequencing short RNA (<200 nt), most protocols call for a coverage ranging from 0.5 to 1.5M reads, under the assumption that discovery rate for new miRNAs is extremely low above 2M reads (42). We argue that the high coverage provided by the TCGA is essential for (i) increasedreliability of miRNA identification (i.e., reporting on expression from a list of 1046 miRNAs); (ii) highlighting non-classical and often low expressing miRNAs.

### Low expressing miRNA act as intrinsic markers for tissue specificity

An unexpected outcome of our study refers to the stability and consistency in the expression of the lowly expressed miRNAs through many samples of the same cancer class. Ample observations show that extreme conditions (e.g., stress, transformation, viral infection, differentiation) lead to drastic changes in miRNA expression profiles in cell types. We postulate that in conditions where miRNA expression patterns dramatically change (e.g., cancer, stem cell differentiation, hypoxia etc.), the tail of the miRNA distribution is hardly affected and consequently cell identity and the origin of the cells remain robust. This interpretation is supported by the gradual manipulating of the data by removal of miRNAs that belong to the lowly expressed tail and assessing the robustness of the classifier to such manipulations (Figure 4).

### miRNA profile can be used for sub-classification of cancer types

TCGA is a fast growing resource and has already reached >10,000 samples which are assigned with 33 cancer types. The methodology presented in this study is applicable for a classification task for any number of diseases’ classes that are supported by enough instances. We showed that the classification performance is very high for a small set of classes (Figure 3B and Table 1). For example, the success rate for Breast Invasive Carcinoma (BRCA) is 96-98%. However, a wide spectrum of survival rates (43,44) is associated with breast cancers, supporting the notion of heterogeneous molecular basis of this disease. Therefore, managing the disease is likely to benefit from a refined classification within the broad class of BRCA. A prognosis profile combining mRNAs, miRNAs and DNA methylation had been proposed (45). In addition to the genes that were implicated as driver mutations, the authors identified number of informative miRNAs such as hsa-miR-328, hsa-miR-484, and hsa-miR-874. While these miRNAs were associated with cell proliferation, angiogenesis and tumor-suppressive functions (46,47), they are poor separators of BRCA from other major tumor types (48). Actually, these miRNAs were implicated in Colon Adenocarcinoma (COAD) (49) and mostly in renal and lung carcinomas (48).

Importantly, each cancer type displays its unique miRNA set. For example, overexpressed miRNAs that were implicated in gastric cancer include miR-17-5p, miR-18a, miR-18b, miR-19a, miR-20b, miR-20a, miR-21, miR-106a, miR-106b, miR-135b, miR-183, miR-340-3p, miR-421 and miR-658. In colon cancer, in addition to the validated oncomiRs, miR-16, miR-31, miR-34a, miR-96, miR-125b and miR-133b were overexpressed. The list of cancer associated miRNAs for pancreatic ductal adenocarcinoma (PDAC) includes miR-15b, miR-95, miR-155, miR-186, miR-190, miR-196a, miR-200b, miR-221 and miR-222 (reviewed (50)). Among the underrepresented miRNAs (called anti-oncomiRs) let-7a-1, miR-143 and miR-145 were validated for a number of cancer types.

We found no statistical evidence for an overlap between the ‘most informative’ miRNAs (e.g., supplemental Table S3, Figure 7B) and miRNAs that are reported in the literature to govern cancer progression. The later includes an extended list of oncomiRs including miR-15, miR-16, miR-17, miR-18, miR-19a, miR-19b, miR-20, miR-21, miR-92, miR-125b, miR-155 and miR-569. We postulate that there is no direct correspondence between the informative tail of lowly expressing miRNAs and the dominating miRNAs that best associated with a transition of a cell to its transformed, cancerous state.

Among the top 20 most informative miRNAs, miR-205 and miR-141 exhibit maximal variability for the 27 classes (by a vector of 81 values, see Materials and Methods). Addition informative miRNAs (Figure 7B) include miR-7-3, miR-31, miR-135a/135b, miR-149, miR-196a-1, miR-200a/200b/200c, miR-215, miR-224, miR-328, miR-429, miR-552, miR-934, miR-944 and miR-1251. In view of the lack of any statistical enrichment between miRNAs assigned as the most significant ones (Figure 7B) and the collection of oncomiRs/ anti-oncomiRs from the literature, we postulate that the multiclass typing and markers for tumorigenesis representing different aspects in the characterization of the disease. Inspecting the top 50 miRNAs from our results addresses (indirectly) a question on the nature of miRNAs that capture most of the classification information. Among the most variable miRNAs, three of the miRNAs (hsa-miR-141, hsa-mir-200a and hsa-mir-200b) belong to one family (mir-8) with a high correlation in their appearance in all 27 classes (r2=0.71-0.74).

### Clinical application and premises

Our results can be used in clinical treatment of patients having cancer of unknown primary origin (CUP). However, toward improvements in clinical guidelines, additional measurements (e.g., immune-histochemical, PCR for mRNA expression) are likely to be beneficial. A set of miRNAs that best classify cancer samples by tissue of origin was presented in view of the success in diagnosis of CUP (20,25,26). A significant overlap between the most informative miRNAs (extracted from (26)) and our SVM protocol was found. By comparing informative 50 candidates from the two studies, we detected 10 shared miRNAs (miR-122, miR-138-1, miR-141, miR-146a, miR-196a, miR-200a, miR-200c, miR-205, miR-31 and miR-9-3). Despite a large difference in the analyzed data (500 versus 8000 samples) and the methodology used (decision tree vs SVM), the consistency among informative miRNAs is intriguing (p-value 2E^5). Ample evidence demonstrated the impact of alteration in hsa-miR-200/hsa-miR-141 expression on proliferation, morphology and aberrant histone acetylation in numerous cancers and cell types. An overview on the contribution of miRNAs for cancer diagnosis, the advantages and limitations of several complementary methods was recently reported (51).

Lastly, we inspected the features that specify the least informative miRNAs in our study. There are 90 miRNAs (out of 1046) that are completely non informative to our task. Namely, in the trichotomy transformation they are uniformly expressed with respect to the 3*27 classes. For these miRNAs the sum of the standard deviations (Table S3) for each of the trichotomy label is zero. Obviously, these miRNAs cannot contribute to the classification task. Interestingly, several validated omcomiRs are among these miRNAs (e.g., miR-16, miR-17, miR-18 and miR-21). This emphasizes the uncoupling between the informative low expressing miRNAs we have identified for the classification task and highly expressed oncomiRs. The level of expression of oncomiRs are used as valuable markers for cell dysregulation and as such are the hallmark of proliferation and transformation in cancer.

## Supplementary information

**Supplementary data Table S1.** The table include two types of information: (i) List of all miRNAs included in the TCGA data; (ii) TCGA ID for each of the samples that were included in the analysis.

**Supplementary data Table S2**. The table depicts the average classification results per class for 100 SVM runs. Numeric source data for Figure 5.

**Supplementary data Table S3.** A list of informative miRNA and their respective ranking. Data source for Figure 7B.

## References

1. Iorio, M.V. and Croce, C.M. (2012) MicroRNA dysregulation in cancer: diagnostics, monitoring and therapeutics. A comprehensive review. EMBO molecular medicine, 4, 143–159.

2. Baffa, R., Fassan, M., Volinia, S., O'Hara, B., Liu, C.G., Palazzo, J.P., Gardiman, M., Rugge, M., Gomella, L.G., Croce, C.M. et al. (2009) MicroRNA expression profiling of human metastatic cancers identifies cancer gene targets. The Journal of pathology, 219, 214–221.

3. Dumortier, O., Hinault, C. and Van Obberghen, E. (2013) MicroRNAs and metabolism crosstalk in energy homeostasis. Cell metabolism, 18, 312–324.

4. Calin, G.A. and Croce, C.M. (2006) MicroRNA signatures in human cancers. Nature reviews. Cancer, 6, 857–866.

5. Volinia, S., Calin, G.A., Liu, C.G., Ambs, S., Cimmino, A., Petrocca, F., Visone, R., Iorio, M., Roldo, C., Ferracin, M. et al. (2006) A microRNA expression signature of human solid tumors defines cancer gene targets. Proceedings of the National Academy of Sciences of the United States of America, 103, 2257–2261.

6. Wang, D., Gu, J., Wang, T. and Ding, Z. (2014) OncomiRDB: a database for the experimentally verified oncogenic and tumor-suppressive microRNAs. Bioinformatics, 30, 2237–2238.

7. Jansson, M.D. and Lund, A.H. (2012) MicroRNA and cancer. Molecular oncology, 6, 590–610.

8. Lujambio, A. and Lowe, S.W. (2012) The microcosmos of cancer. Nature, 482, 347–355.

9. Kitade, Y. and Akao, Y. (2010) MicroRNAs and their therapeutic potential for human diseases: microRNAs, miR-143 and -145, function as anti-oncomirs and the application of chemically modified miR-143 as an anti-cancer drug. Journal of pharmacological sciences, 114, 276–280.

10. Hayes, J., Peruzzi, P.P. and Lawler, S. (2014) MicroRNAs in cancer: biomarkers, functions and therapy. Trends in molecular medicine, 20, 460–469.

11. Lu, J., Getz, G., Miska, E.A., Alvarez-Saavedra, E., Lamb, J., Peck, D., Sweet-Cordero, A., Ebert, B.L., Mak, R.H., Ferrando, A.A. et al. (2005) MicroRNA expression profiles classify human cancers. Nature, 435, 834–838.

12. Di Leva, G. and Croce, C.M. (2013) miRNA profiling of cancer. Current opinion in genetics & development, 23, 3–11.

13. Nishida, N., Yamashita, S., Mimori, K., Sudo, T., Tanaka, F., Shibata, K., Yamamoto, H., Ishii, H., Doki, Y. and Mori, M. (2012) MicroRNA-10b is a prognostic indicator in colorectal cancer and confers resistance to the chemotherapeutic agent 5-fluorouracil in colorectal cancer cells. Annals of surgical oncology, 19, 3065–3071.

14. Bockhorn, J., Dalton, R., Nwachukwu, C., Huang, S., Prat, A., Yee, K., Chang, Y.F., Huo, D., Wen, Y., Swanson, K.E. et al. (2013) MicroRNA-30c inhibits human breast tumour chemotherapy resistance by regulating TWF1 and IL-11. Nature communications, 4, 1393.

15. Bryant, R.J., Pawlowski, T., Catto, J.W., Marsden, G., Vessella, R.L., Rhees, B., Kuslich, C., Visakorpi, T. and Hamdy, F.C. (2012) Changes in circulating microRNA levels associated with prostate cancer. British journal of cancer, 106, 768–774.

16. Pentheroudakis, G., Briasoulis, E. and Pavlidis, N. (2007) Cancer of unknown primary site: missing primary or missing biology? The oncologist, 12, 418–425.

17. Pavlidis, N. and Pentheroudakis, G. (2010) Cancer of unknown primary site: 20 questions to be answered. Annals of oncology : official journal of the European Society for Medical Oncology/ESMO, 21 Suppl 7, vii303–307.

18. Fizazi, K., Greco, F.A., Pavlidis, N., Daugaard, G., Oien, K., Pentheroudakis, G. and Committee, E.G. (2015) Cancers of unknown primary site: ESMO Clinical Practice Guidelines for diagnosis, treatment and follow-up. Annals of oncology : official journal of the European Society for Medical Oncology /ESMO, 26 Suppl 5, v133–138.

19. Greco, F.A., Oien, K., Erlander, M., Osborne, R., Varadhachary, G., Bridgewater, J., Cohen, D. and Wasan, H. (2012) Cancer of unknown primary: progress in the search for improved and rapid diagnosis leading toward superior patient outcomes. Annals of oncology : official journal of the European Society for Medical Oncology / ESMO, 23, 298–304.

20. Rosenwald, S., Gilad, S., Benjamin, S., Lebanony, D., Dromi, N., Faerman, A., Benjamin, H., Tamir, R., Ezagouri, M., Goren, E. et al. (2010) Validation of a microRNA-based qRT-PCRtest for accurate identification of tumor tissue origin. Modern pathology : an official journal of the United States and Canadian Academy of Pathology, Inc, 23, 814–823.

21. Varadhachary, G.R., Spector, Y., Abbruzzese, J.L., Rosenwald, S., Wang, H., Aharonov, R., Carlson, H.R., Cohen, D., Karanth, S., Macinskas, J. et al. (2011) Prospective gene signature study using microRNA to identify the tissue of origin in patients with carcinoma of unknown primary. Clinical cancer research : an official journal of the American Association for Cancer Research, 17, 4063–4070.

22. Bishop, J.A., Benjamin, H., Cholakh, H., Chajut, A., Clark, D.P. and Westra, W.H. (2010) Accurate classification of non-small cell lung carcinoma using a novel microRNA-based approach. Clinical cancer research : an official journal of the American Association for Cancer Research, 16, 610–619.

23. Gilad, S., Lithwick-Yanai, G., Barshack, I., Benjamin, S., Krivitsky, I., Edmonston, T.B., Bibbo, M., Thurm, C., Horowitz, L., Huang, Y. et al. (2012) Classification of the four main types of lung cancer using a microRNA-based diagnostic assay. The Journal of molecular diagnostics : JMD, 14, 510–517.

24. Ferracin, M., Pedriali, M., Veronese, A., Zagatti, B., Gafa, R., Magri, E., Lunardi, M., Munerato, G., Querzoli, G., Maestri, I. et al. (2011) MicroRNA profiling for the identification of cancers with unknown primary tissue-of-origin. The Journal of pathology, 225, 43–53.

25. Meiri, E., Mueller, W.C., Rosenwald, S., Zepeniuk, M., Klinke, E., Edmonston, T.B., Werner, M., Lass, U., Barshack, I., Feinmesser, M. et al. (2012) A second-generation microRNA-based assay for diagnosing tumor tissue origin. The oncologist, 17, 801–812.

26. Rosenfeld, N., Aharonov, R., Meiri, E., Rosenwald, S., Spector, Y., Zepeniuk, M., Benjamin, H., Shabes, N., Tabak, S., Levy, A. et al. (2008) MicroRNAs accurately identify cancer tissue origin. Nature biotechnology, 26, 462–469.

27. De Cecco, L., Dugo, M., Canevari, S., Daidone, M.G. and Callari, M. (2013) Measuring microRNA expression levels in oncology: from samples to data analysis. Critical reviews in oncogenesis, 18, 273–287.

28. Kosaka, N., Iguchi, H. and Ochiya, T. (2010) Circulating microRNA in body fluid: a new potential biomarker for cancer diagnosis and prognosis. Cancer science, 101, 2087–2092.

29. Fesler, A., Jiang, J., Zhai, H. and Ju, J. (2014) Circulating microRNA testing for the early diagnosis and follow-up of colorectal cancer patients. Molecular diagnosis & therapy, 18, 303–308.

30. Monzon, F.A. and Koen, T.J. (2010) Diagnosis of metastatic neoplasms: molecular approaches for identification of tissue of origin. Archives of pathology & laboratory medicine, 134, 216–224.

31. Gore, J., Craven, K.E., Wilson, J.L., Cote, G.A., Cheng, M., Nguyen, H.V., Cramer, H.M., Sherman, S. and Korc, M. (2015) TCGA data and patient-derived orthotopic xenografts highlight pancreatic cancer-associated angiogenesis. Oncotarget, 6, 7504–7521.

32. Zhang, W. (2014) TCGA divides gastric cancer into four molecular subtypes: implications for individualized therapeutics. Chinese journal of cancer, 33, 469–470.

33. Ho, D.W., Kai, A.K. and Ng, I.O. (2015) TCGA whole-transcriptome sequencing data reveals significantly dysregulated genes and signaling pathways in hepatocellular carcinoma. Frontiers of medicine, 9, 322–330.

34. Zhu, Y., Qiu, P. and Ji, Y. (2014) TCGA-assembler: open-source software for retrieving and processing TCGA data. Nature methods, 11, 599–600.

35. Chang, C.C., Hsu, C.W. and Lin, C.J. (2000) The analysis of decomposition methods for support vector machines. IEEE transactions on neural networks / a publication of the IEEE Neural Networks Council, 11, 1003–1008.

36. Abraham, A., Pedregosa, F., Eickenberg, M., Gervais, P., Mueller, A., Kossaifi, J., Gramfort, A., Thirion, B. and Varoquaux, G. (2014) Machine learning for neuroimaging with scikit-learn. Frontiers in neuroinformatics, 8, 14.

37. Bosson, A.D., Zamudio, J.R. and Sharp, P.A. (2014) Endogenous miRNA and target concentrations determine susceptibility to potential ceRNA competition. Molecular cell, 56, 347–359.

38. Jackson, A.L., Burchard, J., Schelter, J., Chau, B.N., Cleary, M., Lim, L. and Linsley, P.S. (2006) Widespread siRNA “off-target” transcript silencing mediated by seed region sequence complementarity. Rna, 12, 1179–1187.

39. Tay, Y., Rinn, J. and Pandolfi, P.P. (2014) The multilayered complexity of ceRNA crosstalk and competition. Nature, 505, 344–352.

40. Hansen, T.B., Jensen, T.I., Clausen, B.H., Bramsen, J.B., Finsen, B., Damgaard, C.K. and Kjems, J. (2013) Natural RNA circles function as efficient microRNA sponges. Nature, 495, 384–388.

41. Balaga, O., Friedman, Y. and Linial, M. (2012) Toward a combinatorial nature of microRNA regulation in human cells. Nucleic acids research, 40, 9404–9416.

42. Metpally, R.P., Nasser, S., Malenica, I., Courtright, A., Carlson, E., Ghaffari, L., Villa, S., Tembe, W. and Van Keuren-Jensen, K. (2013) Comparison of Analysis Tools for miRNA High Throughput Sequencing Using Nerve Crush as a Model. Frontiers in genetics, 4, 20.

43. Cancer Genome Atlas, N. (2012) Comprehensive molecular portraits of human breast tumours. Nature, 490, 61–70.

44. Tahiri, A., Leivonen, S.K., Luders, T., Steinfeld, I., Ragle Aure, M., Geisler, J., Makela, R., Nord, S., Riis, M.L., Yakhini, Z. et al. (2014) Deregulation of cancer-related miRNAs is a common event in both benign and malignant human breast tumors. Carcinogenesis, 35, 76–85.

45. Volinia, S. and Croce, C.M. (2013) Prognostic microRNA/mRNA signature from the integrated analysis of patients with invasive breast cancer. Proceedings of the National Academy of Sciences of the United States of America, 110, 7413–7417.

46. Dai, Y., Sui, W., Lan, H., Yan, Q., Huang, H. and Huang, Y. (2009) Comprehensive analysis of microRNA expression patterns in renal biopsies of lupus nephritis patients. Rheumatology international, 29, 749–754.

47. Nohata, N., Hanazawa, T., Kikkawa, N., Sakurai, D., Fujimura, L., Chiyomaru, T., Kawakami, K., Yoshino, H., Enokida, H., Nakagawa, M. et al. (2011) Tumour suppressive microRNA-874 regulates novel cancer networks in maxillary sinus squamous cell carcinoma. British journal of cancer, 105, 833–841.

48. Leidinger, P., Keller, A., Borries, A., Huwer, H., Rohling, M., Huebers, J., Lenhof, H.P. and Meese, E. (2011) Specific peripheral miRNA profiles for distinguishing lung cancer from COPD. Lung cancer, 74, 41–47.

49. Li, H.G., Zhao, L.H., Bao, X.B., Sun, P.C. and Zhai, B.P. (2014) Meta-analysis of the differentially expressed colorectal cancer-related microRNA expression profiles. European review for medical and pharmacological sciences, 18, 2048–2057.

50. Paranjape, T., Slack, F.J. and Weidhaas, J.B. (2009) MicroRNAs: tools for cancer diagnostics. Gut, 58, 1546–1554.

51. Pichler, M. and Calin, G.A. (2015) MicroRNAs in cancer: from developmental genes in worms to their clinical application in patients. British journal of cancer, 113, 569–573.

